# Focused ultrasound enhanced antibody delivery for the treatment of Parkinson’s Disease

**DOI:** 10.1101/2024.09.13.611071

**Authors:** Nancy Kwon, Alec J. Batts, Hairong Zhang, Carlos Sierra Sanchez, Vernice R. Jackson-Lewis, Serge Przedborski, Elisa E. Konofagou

## Abstract

Treatment of neurological disorders is partly impeded by the size of large pharmacological agents which are thereby unable to bypass the blood-brain barrier (BBB). Focused ultrasound (FUS) in conjunction with systemically administered microbubbles has been shown to safely, non-invasively and transiently open the BBB, allowing the passage of large biomolecules to the brain parenchyma through the otherwise impermeable barrier. This pilot study assessed the feasibility of FUS-mediated delivery of an anti-alpha-synuclein (α-syn) monoclonal antibody (mAb) in Parkinson’s disease (PD) mouse models that exhibit α-syn aggregates. Mice (n=21) underwent FUS on a weekly basis over the course of 2-3 weeks, followed by a one-month survival period. MRI and microscopy were performed to confirm BBB opening with FUS and visualize antibody delivery. Safety was assessed in vivo using passive cavitation detection and immunohistochemistry to evaluate microglial and astrocyte activity ex vivo. It was found that treatment sessions for multiple FUS sessions of targeted antibody delivery was feasible in alpha-synuclein models facilitating immunotherapeutics for PD.

## INTRODUCTION

Parkinson’s Disease (PD) is expected to be one of the fastest growing neurological disorders in prevalence, disability, and death based on the global burden of disease study which forecasts the incidence of PD to increase by 21.7% when taking into account the trajectory of the aging population (1). While the gold standard of PD treatment has focused on symptom management by treating dopaminergic loss, other treatments currently under investigation have focused on disease modification by targeting pathophysiology with cell-based therapeutics, gene therapy with viral vectors, and more recently, immunotherapy (2–5). Immunotherapy for neurodegenerative diseases such as Alzheimer’s disease has shown modest success, targeting pathological amyloid-beta plaques with recent FDA approvals for lecanemab, aducanumab, and donanemab. Immunization strategies in PD have targeted against α-synuclein (αsyn), a major component of Lewy Bodies considered a hallmark of PD (PMID: 12971891). Preclinical antibody studies in PD mouse models expressing pathological α-syn, have shown a reduction in α-syn burden coinciding with improvement in motor and cognitive performance (6–10). Given these preclinical studies, clinical trials have been performed revealing that the humanized anti-α-syn antibodies and regimens were well-tolerated and markedly reduce circulating α-syn levels [29, 30]. Despite these encouraging human data, two phase 2 trials examining monoclonal antibodies treatment of PD failed to show significant benefits [31, 32]. However, neither trial demonstrated appropriate target engagement in the central nervous system (CNS), which we consider a critical flaw as most antibodies do not permeate the BBB or plasma membranes [33]. Therefore, we posit that more robust neuroprotection requires higher levels of anti–α-syn antibody accumulation in the brain, particularly in those regions prone to α-syn pathology.

Drug therapeutics for the treatment of CNS diseases have largely been impeded by the presence of the BBB, preventing molecules larger than 400-600 Da from diffusing across the barrier (13). However, focused ultrasound (FUS) coupled with systemically administered microbubbles (MBs) has been shown to provide a promising therapeutic alternative for targeted CNS delivery noninvasively, transiently, and directly to the brain parenchyma while leaving other regions unaffected (14–16,16–23).

FUS-BBBO allows for the non-invasive, localized drug delivery with target specificity that allows for penetration of deeper brain regions. Previous publications have shown that FUS allows for the passage of large molecules such as neurotrophic factors (12-13 kDa) and viral vectors (∼60-90 kDa) to specific brain regions (22,24,25). Furthermore, the effects of FUS-BBBO alone indicate a reduction of amyloid plaques and tau in animal models (17,26–28). Our group has demonstrated FUS-BBBO mediated delivery of gene vectors and proteins resulted in 76% and 52% higher terminal and dendrite density (respectively) than in untreated animals after FUS-mediated delivery (17).

The objective of this study is to demonstrate the feasibility of using FUS-BBBO for CNS passive immunotherapy for neurodegenerative disorders such as PD. We hereby show that FUS-BBBO anti αsyn antibody delivery in old and young mice with pathological αsyn aggregates that were treated in multiple sessions on a weekly basis.

## MATERIALS AND METHODS

### Animals

Procedures described herein were approved by and performed in accordance with guidelines set by Columbia University’s Institutional Animal Care and Use Committee (IACUC). Transgenic mice expressing αsyn mutations were used for the pilot and long-term studies. These mutations express αsyn aggregates which genetic studies have found to be a primary component of Lewy bodies seen in familial cases of PD (29,30). Mice bearing the A30P mutation (12-15 months) were used in the pilot study to confirm FUS-mediated mAB delivery and 18 mice were used in the long-term study; 6 mice expressing the SNCA or A30P mutation (12-15 months) and 12 A53TmVLEM83 male mice that express A53T mutations (6-7 months).

### Focused ultrasound with passive cavitation detection (PCD)

As previously described (22,31), a single-element, concave transducer with a center frequency of 1.5 MHz and focal depth of 60 mm (outer/inner radius: 30 mm/11.2 mm) was driven by a function generator (Agilent, Palo Alto, CA, USA) and amplified with a 50-dB power amplifier (ENI Inc., Rochester, NY, USA) (Figure 1). Concurrently, PCD was performed with the pulse-echo transducer (center frequency: 7.5 MHz, focal length: 60 mm, Olympus NDT, Waltham, MA, USA) that is confocally aligned with the FUS transducer to acquire signals from oscillating MB when the FUS transducer is in transmit mode. This is connected to a digitizer (Gage Applied Technologies, Inc., Lachine, QC, Canada) with a pulse-receiver that has a 20 dB-amplification (Part No. 5072; Olympus Industrial, Waltham, MA, USA). A cone with an opening covered by translucent film is fitted to the FUS transducer and filled with degassed water and mounted onto the system. The following FUS parameters were applied at each region of interest (ROI): 60 second pulsed sequence of 10,000 cycles, pulse repetition frequency (PRF) of 5 Hz, and an estimated derated peak negative pressure (PNP) of .450 KPa, with 18% attenuation (17,18).

**Figure 1.**
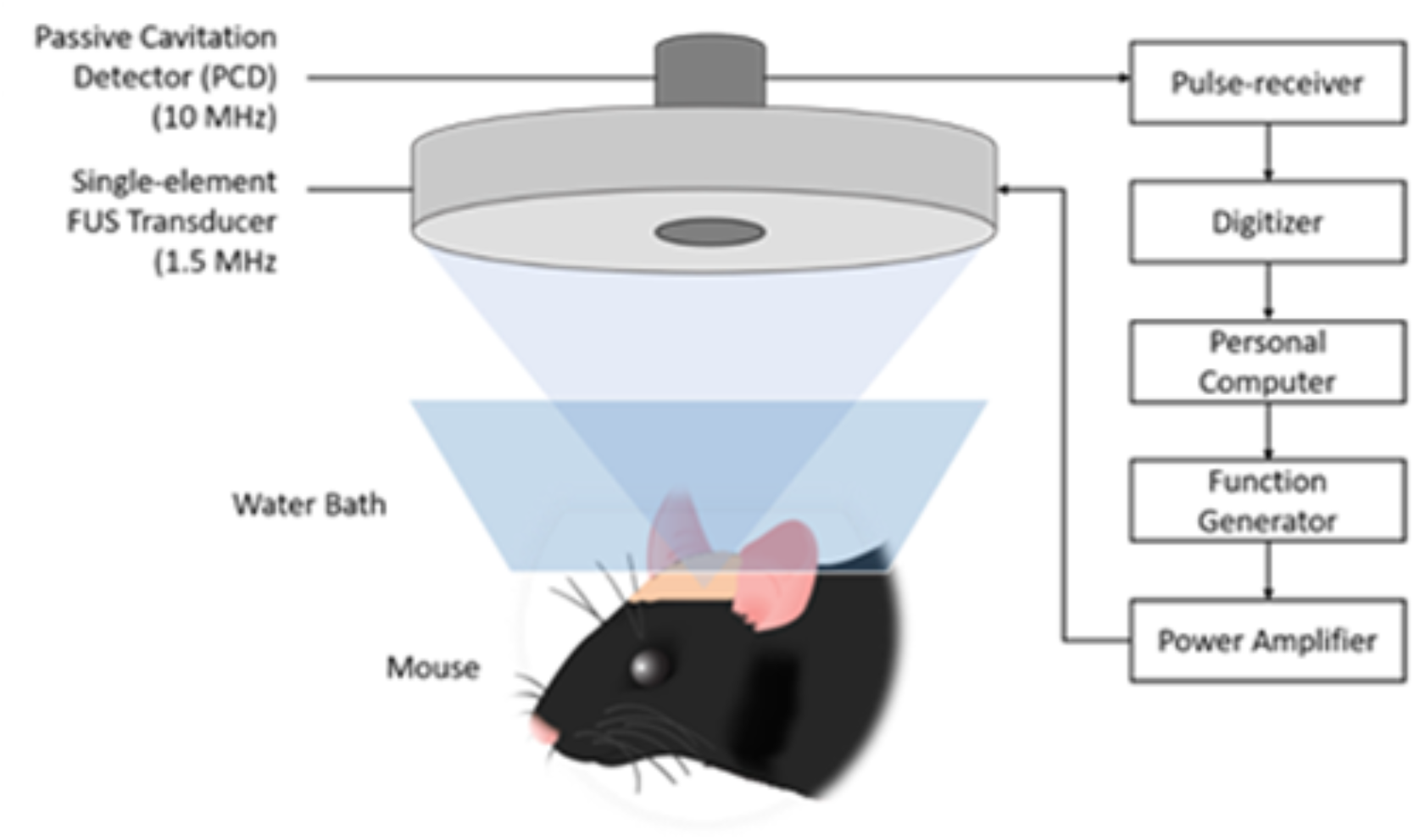
Focused ultrasound (FUS) system with single-element transducer and passive cavitation detection (PCD).

### Magnetic Resonance Imaging (MRI)

To confirm BBB opening, mice underwent a contrast-enhanced MRI in a 9.4 T vertical bore system (Bruker, Boston, MA). Shortly after FUS-BBBO, Mice were injected intraperitoneally with gadolinium then imaged using a T1-weighted 2D FLASH sequence (TR/TE 230/3.3 ms, flip angle 70, resolution 100 μm × 100 μm × 400 μm). A T2-weighted Turbo RARE sequence (TR/TE 2500/30 ms, echo spacing 10 ms, resolution 100 × 100 × 400 μm) was also acquired to evaluate safety by the absence of edema or hemorrhages.

### Microbubbles

Size-isolated, lipid-coated microbubbles (MB) were formed as previously reported (32). Briefly, two lipid compounds (1,2-distearoyl-sn-glycero-3-phospho-choline (DSPC) and 1,2-distearoyl-sn-glycero-3-phosphoethanolamine-N-[methoxy(polyethylene glycol)-2000] (DSPC-mPEG2000; Avanti Polar Lipids Inc., Alabaster, AL, USA) were dissolved in a solution comprised of 10% glycerin, 10% propylene glycol, and 80% phosphate-buffered saline (PBS) and alternated between an ultrasonic warm water bath and sonication with a high-power sonication tip (Branson; Brookfield, CT) until the final solution was clear. The solution was allowed to cool then exposed to perfluorobutane gas (FluoroMed L.P., Round Rock, TX, USA) forming the microbubbles, then size-isolated in a series of alternating high and low centrifuge cycles (32). Final concentration was measured with a multisizer (Beckman Coulter; Pasedena, CA) to confirm mean diameter and concentration.

### Monoclonal antibody (mAB)

A rabbit anti-αsyn antibody conjugated with Alexa Fluor 488 (mAB-AF488) (ab195025, MJFR1; Abcam, MA, USA) and an untagged version of the mAB were used for the pilot and long-term study. This antibody was selected due to strong immunoreactivity as defined by measuring intensity of DAB-IHC observed on immunohistochemistry (IHC) of brain sections in transgenic mice previously performed by our collaborator.

### Study Design

A pilot study involved A30P and SNCA transgenic mice, 12-15 months of male and female mice, to confirm targeted antibody delivery with FUS in the subcortical brain regions of the caudate putamen (CPu) and substantia nigra (SN). Mice underwent FUS-mAB, were then sacrificed by transcardial perfusion 3-5 hours after FUS delivery of the mAB for tissue to be collected and processed for microscopy to detect fluorescently tagged mAB in brain tissue. Subsequently, a small-scale survival study was conducted with old and young transgenic mice to evaluate the feasibility of a long-term ‘treatment’ schema as shown in Figure 2. This cohort received FUS-mAB at three separate sessions a week apart, then survived for one month before sacrifice.

**Figure 2.**
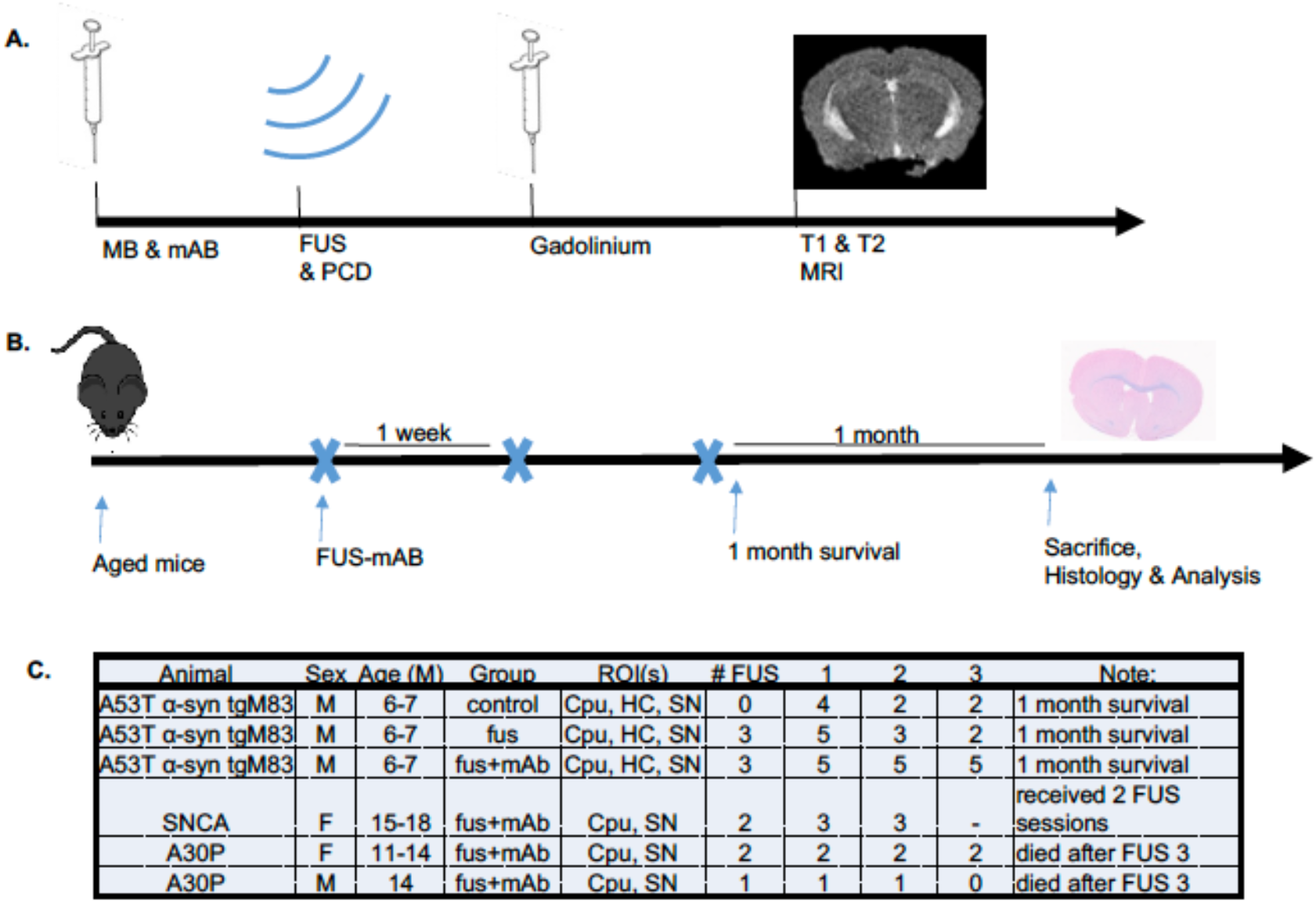
(A) Schema of FUS session, (B) long-term safety evaluation, and (C) corresponding chart of animals involved in the study along with a survival profile. (C) Survival of transgenic (Tg) mice in long-term study with survival numbers after each FUS session. Young Tg+ mice (6-7 months) that received FUS and antibody (mAB) survived after three FUS sessions and a 1-month survival period, whereas only half the control group survived the same length of time. In an ‘older’ cohort, SNCA mice only received two completed FUS-mAB sessions after early death in A30P shortly after completing a third session.

During the FUS session, the mouse was anesthetized with isoflurane and oxygen that was delivered to a anesthesia mask fitted for rodent and placed prone throughout the procedure. The animal’s head was fixed in a stereotaxic frame (Kopf Instruments; Tujunga, CA), with the FUS system positioned overhead. Prep involved fur removal and ultrasound gel applied to couple the transducer surface to the scalp. A 27g needle (Terumo; Somerset, NJ) was inserted into the tail for intravenous (IV) injections of MB and mAB.

Each FUS-session involved a bolus injection of MB (50 µL; 8×108 MB/mL) and mAB (50 µL; 1 mg/mL) diluted in saline before FUS-mAB. Regions were targeted with FUS following coordinates relative to lambda as determined from a mouse brain atlas (33): CPu: -1.5 mm lateral, +2.5 mm anterior, HC: -2.5 lateral, +1 anterior, and SN: -2 mm lateral, -1.5 mm posterior (33). After each FUS session on the same day, an MRI was conducted to confirm opening and safety.

Confirmation of mAB delivery was observed in the CPu at 50 µg and 100 µg doses (Figure 3). Subsequent studies delivered 50 µg of mAB to the CPu, SN, and HC at each FUS session followed by a 1-month survival period. Following FUS, animals were sacrificed by the methods as previously reported. Briefly, animals were anesthetized then euthanized via transcardial perfusion using 60 mL of 4% paraformaldehyde (PFA) and 30 mL pharmacological grade 1 x phosphate buffered saline (PBS), then brain tissue was excised and prepared for microscopy (17,17,18). Floating sections were collected at 35 µm then immunostained for αsynuclein, 1:500 (R&D Systems; Minneapolis, MN), microglia with Iba-1, 1:500 (ionized calcium-binding adaptor molecule-1; Abcam, Cambridge, MA), and astrocytes with GFAP, 1:500 (glial-fibrillary actin protein; Abcam, Cambridge, MA). Images were acquired at 20x with a confocal microscope (Nikon, Tokyo, Japan) or a Zeiss LSM-700 (Oberkochen, Germany).

**Figure 3.**
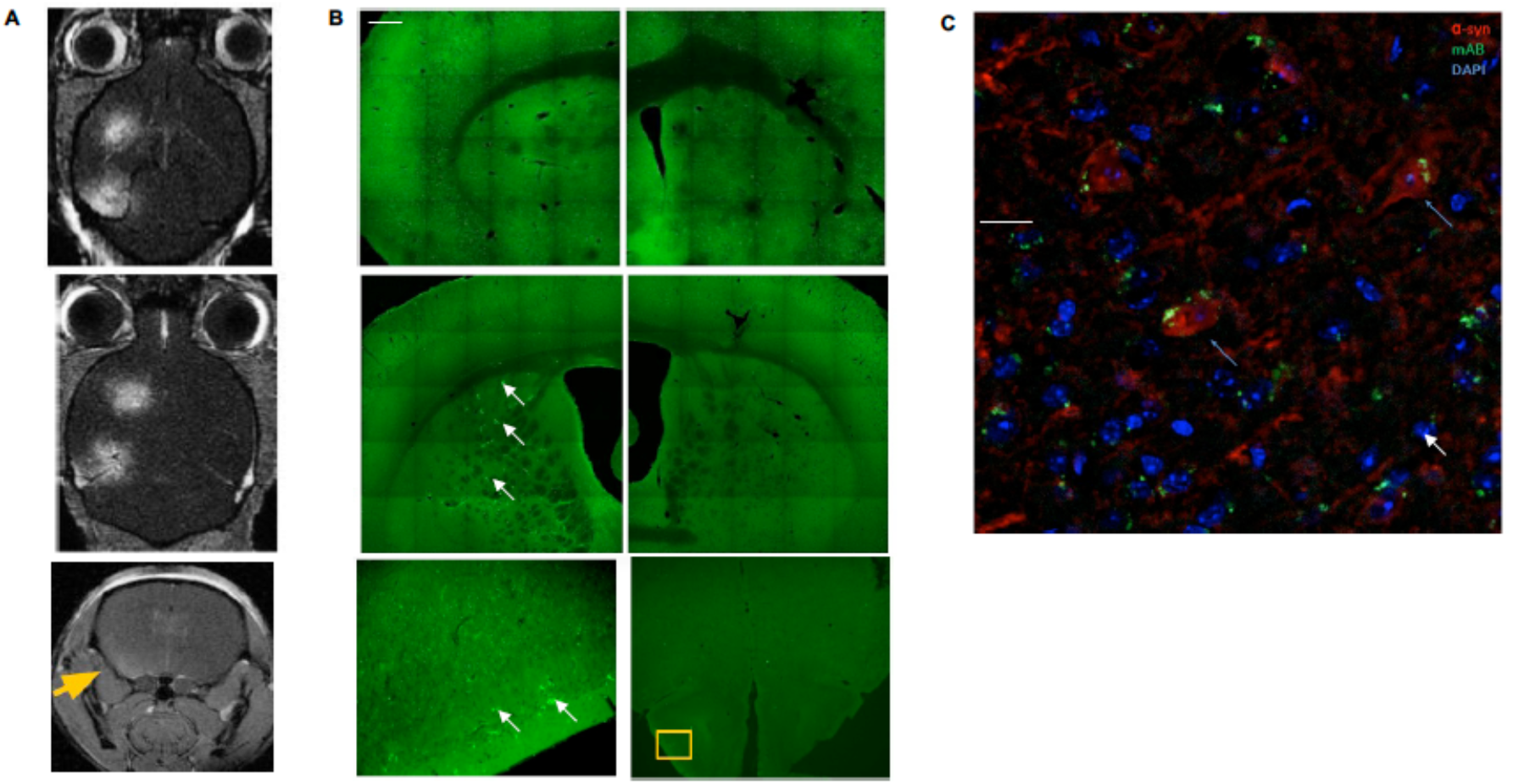
FUS-mAB confirmation compared to the contralateral side. T1-weighted contrast enhanced MRI (Left column) confirming FUS-BBBO and corresponding microscopy mages (middle) compare left (treated) and right (untreated) hemispheres at 35 µg (A), 50 µg (B), 100 µg (C). (D) Colocalization of mAB-AF488 (green) and alpha-synuclein (red) with DAPI after FUS-mAB delivery.

## RESULTS DISCUSSION

Safety and reproducibility of FUS-mAB was evaluated in a long-term study with MRI, PCD, and histology (Figure 4). T2-weighted scans showed no hyperintense voxels indicative of damage after each session. PCD analysis of stable and inertial cavitation doses from each ROI after each FUS session was used to evaluate BBBO based on previous studies corroborating cavitation dose with BBBO at 450 kPa which was also observed in this study, demonstrating consistent FUS-BBBO with no significant difference in SCD (p<0.05) but significant difference in the ICD (p<0.01), confirmed by one-way ANOVA and Bonferroni post-hoc multiple comparisons test (18,31,34).

**Figure 4.**
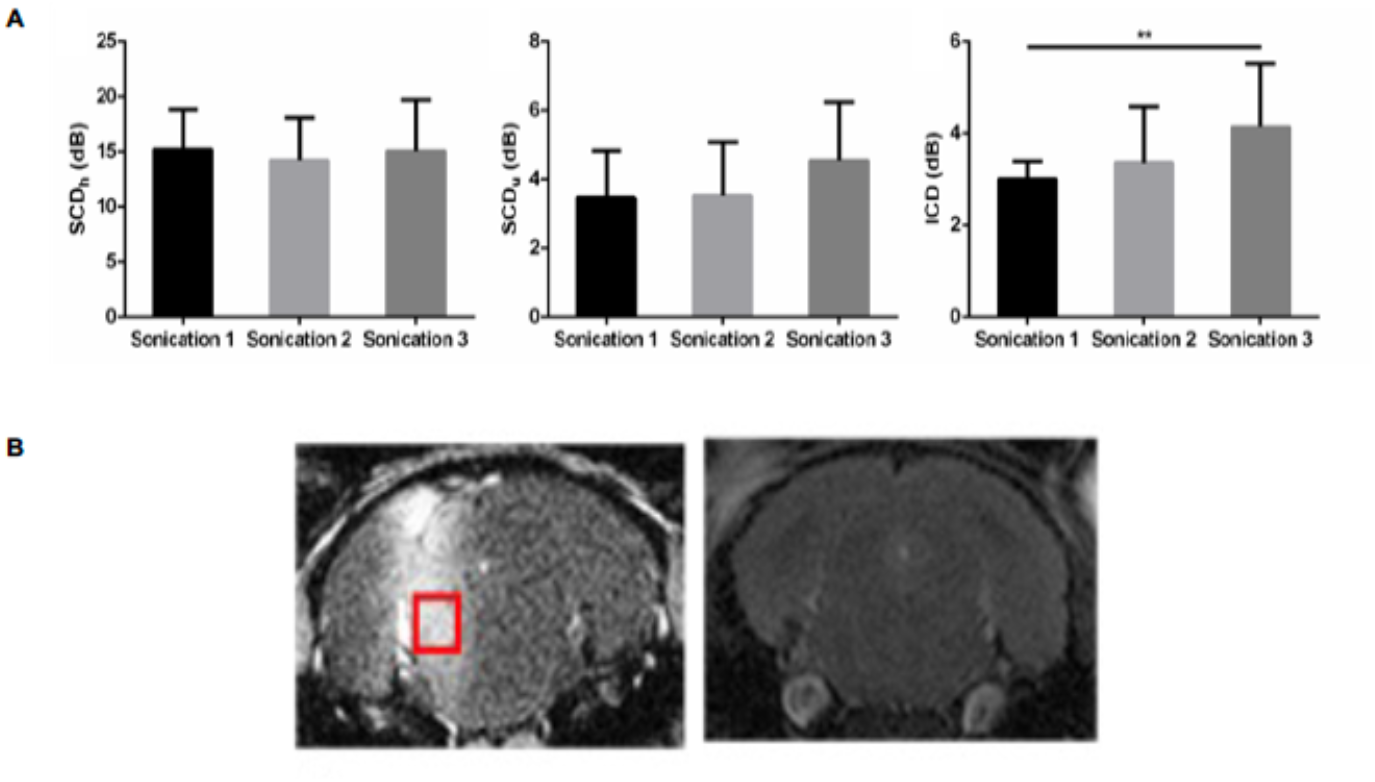
Safety and reproducibility of FUS evaluated with PCD (A) and T2-weighted MRI scans (B). Safety analysis based on cavitation signals show no significant difference in stable cavitation (p>0.05). However, a significant difference of inertial cavitation (p<0.01), confirmed by one-way ANOVA followed by a Bonferroni post-hoc multiple comparisons test. (B) T1- and T2-weighted MRI images of FUS-BBBO in the SN which confirmed BBBO and absence of edema after FUS session.

Histology was performed to assess microgliosis (Iba-1) and phagocytic activity (Cd68), in addition to neuronal cell viability (NEUN) after the study period and further analysis conducted using Image J to preliminary analyze immunofluorescent intensity between control and FUS-mAB revealed no significant difference in phagocytic activity between the FUS-treated and control group which demonstrates the long-term safety of FUS-mediated drug delivery (Figure 5). Moreover, no significant changes in microglia morphology were detected qualitatively. Analysis of neuronal cells also show no significant difference after FUS-mAB.

**Figure 5.**
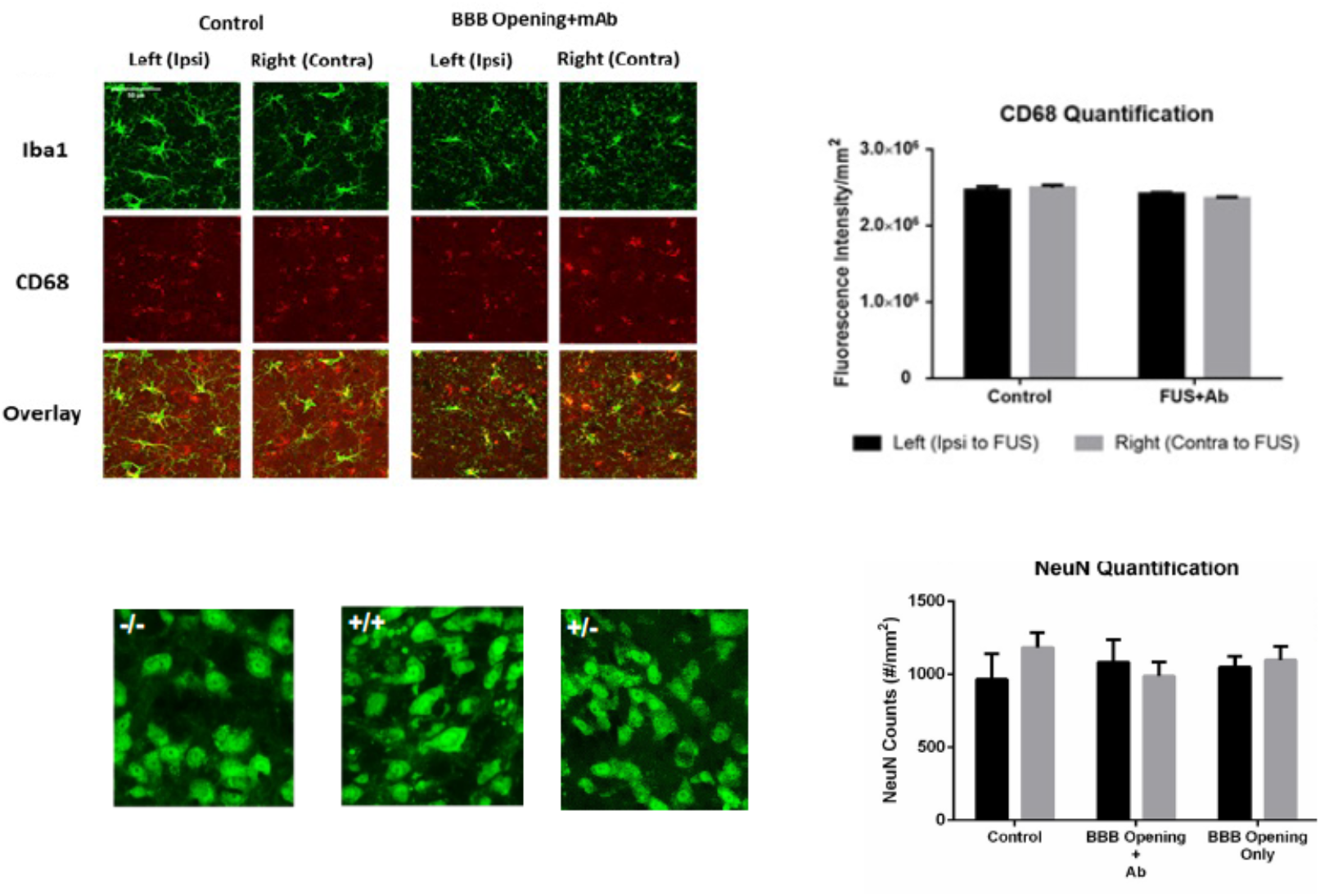
Safety further evaluated 1 month after last FUS-mAB session in A53T mice. (TOP) Microglia and associated phagocytic activity as observed with histology. Intensity quantification did not show significant altered activity between FUS-mAB and control A53T mice in the long-term study. Scale bars denote 50 µm. (BOTTOM) Quantification of Neun intensity show no difference between control, BBB opening with and without mAB.

Our findings presented herein indicate that FUS-mAB delivery is a feasible technique that could be safely implemented in a clinical setting; potentially with a similar treatment schema involving multiple targeted regions in a single treatment with the capability of repeated drug administration if deemed necessary. Future directions will focus on optimizing the antibody-delivery treatment protocol in addition to assessing the pharmacokinetics and pharmacodynamics of antibody delivery to the brain parenchyma to elicit a successful immune response. Localized delivery directly in the brain parenchyma may also reduce side effects often experienced through systemic routes such as oral medications which are not targeted and require the drug to be broken down by the gastrointestinal system.

More studies in our field are needed to fully understand the immune response of FUS-BBBO and of targeted immunotherapies to properly assess feasibility of clinical translation. Our study demonstrated FUS-mediated antibody delivery can be performed safely and repeatable delivery into affected PD regions of the caudate putamen of mice.

## CONCLUSION

The study presented herein demonstrates the utility of FUS-therapeutics in the treatment of PD on a weekly schedule, in the absence of deleterious effects. We showed that FUS parameters used in the current study are below the threshold for damage as shown by MRI scans and post-mortem histology that corroborate our findings on the safety of FUS during the active treatment phase and long-term survival.

## ACKNOWLEDGEMENTS

This study was funded by the National Institutes of Health (EB009041, AG038961, NS111176, AG064596), and the Focused Ultrasound Foundation.

The authors also would wish to acknowledge the Confocal and Specialized Microscopy Shared Resources (CSMSR) of the Herbert Irving Comprehensive Cancer Center (HICCC) of Columbia University, supported by NIH grant #P30 CA013696.

We would also wish to acknowledge Pablo Abreu for his administrative support, Marilena Karakatsani, Robin Ji, and Antonis Pouliopolous for their assistance with data and analysis.

